# Force generation in the coiling tendrils of *Passiflora caerulea*

**DOI:** 10.1101/2023.02.08.527445

**Authors:** Frederike Klimm, Thomas Speck, Marc Thielen

## Abstract

Tendrils of climbing plants coil along their length and thus form a striking helical spring and generate tensional forces. We have found that, for tendrils of the passion flower *Passiflora caerulea*, the generated force lies in the range of 6-140 mN, which is sufficient to lash the plant tightly to its substrate. Further, we revealed that the generated force strongly correlates with the water status of the plant. By combining force measurements with anatomical investigations and dehydration-rehydration experiments on both entire tendril segments and isolated lignified tissues, we are able to propose a two-phasic principle of spring formation: First, during the free coiling phase, the tendril coiling is based on the active contraction of a fiber ribbon in interaction with the surrounding parenchyma as resistance layer. Second, in a stabilization phase, the entire center of the coiled tendril lignifies, stiffening the spring and securing its function independent of hydration status.

## Introduction

Plants as rooted sessile organisms are not usually associated with movement. They do however move substantially, as is particularly evident in climbing plants (e.g., [1]). Instead of producing a stiff self-supporting trunk, these plants rely on supporting host structures to reach upwards to the light (e.g., [2], [3]) (fig. 1 a). They attach to their supports, for instance, by tendrils. These filamentary organs are sensitive to contact [1] and either possess adhesive discs at their tips, as in, for example, the tendrils of *Parthenocissus tricuspidata* [4] or *Passiflora discophora* [5], or “grasp” slender supports by coiling around them (a process termed as contact coiling, e.g., [6]). Eventually, the tendrils coil along their length axis in a movement referred to as free coiling (e.g., [6]). They do so by developing a length difference between their prospective convex and concave sides (e.g., [7]), thereby creating an intrinsic curvature and eventually resulting in helical coiling. Since the tendril’s ends are fixed in position, free coiling results in a particular spring-like shape with a minimum of one perversion, which links two helices of opposite handedness [8], [9] (fig. 1 b).

**Fig. 1:**
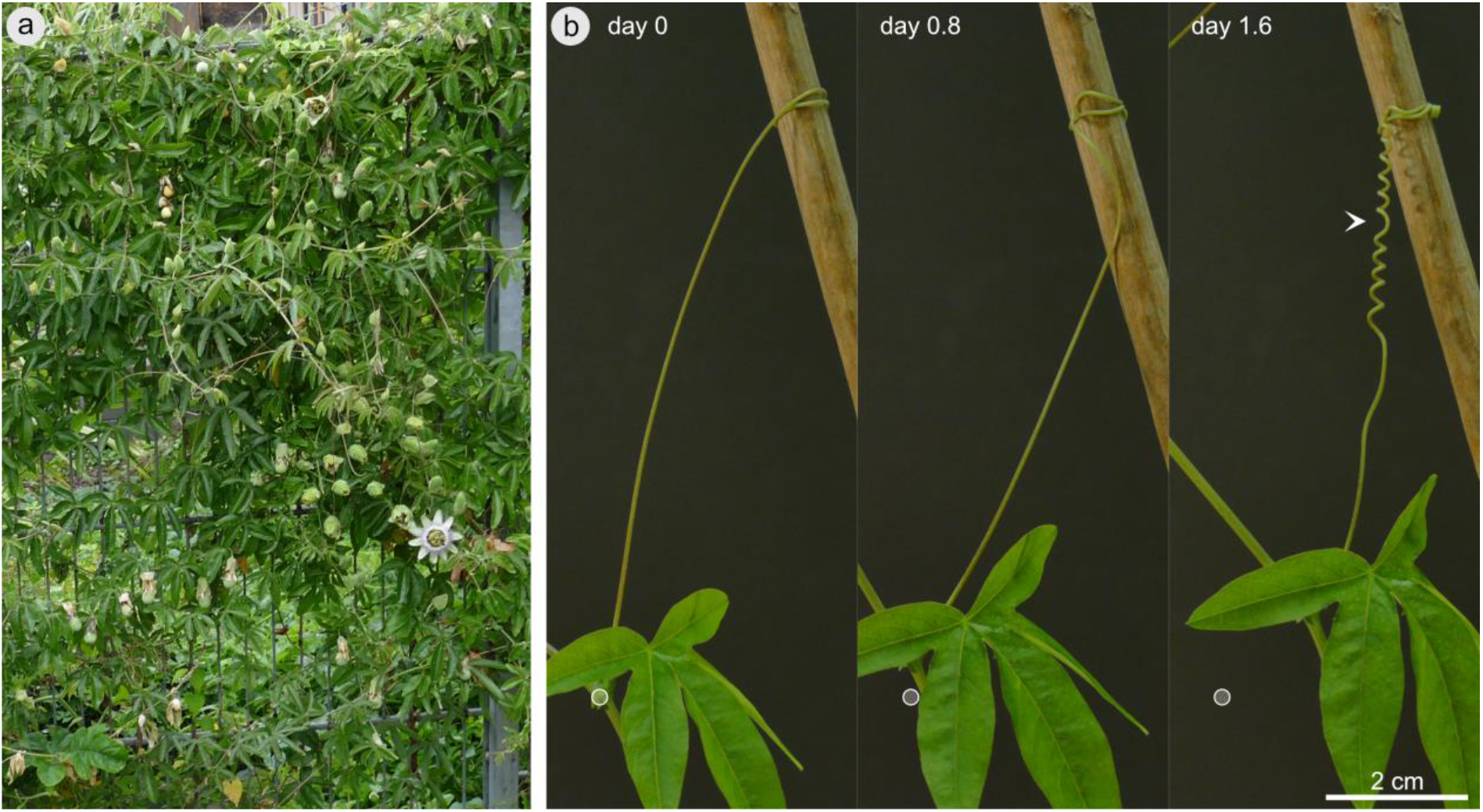
Climbing passion flower *Passiflora caerulea*. Habitus of *P. caerulea* climbing on a metal fence in the outdoor area of the Botanic Garden of the University of Freiburg **(a)**. Images of a free-coiling *P. caerulea* tendril lashing the plant stem closer to its support (white circle indicates reference position of the node at day 0) and forms a spring-like shape consisting of two helices of opposite handedness connected by an inversion point (perversion, arrowhead) **(b)**. See also time-lapse supplement video 1.

Several mechanisms have been suggested to cause the length difference leading to tendril curvature and eventually to free or contact coiling, including differential growth, turgor changes [1], [10], [11], [6], and the contraction of G-fibers [12]. G-fibers are fiber cells that, in addition to having the typical primary and secondary cell wall layers, possess a gelatinous-or G-layer that enables them to contract (e.g., [13], [14]). A prominent instance of the occurrence of these fibers is the tension wood in trees (e.g., [15], [14]) and contractile roots [16]. As mentioned above, this type of fiber is also associated with twining and coiling in many climbing plant species [12], [13], [17], [18], [19]. However, the correct identification of the G-layer in some of the cited works has been brought into question [13]. In tendrils, the presence of contractile G-fibers suggests a straightforward mechanism for bending and thus coiling, notably by means of a bilayer that combines one contractile layer with a non-or, at least, a less contractile antagonistic layer. Further, the contractile force of G-fibers is suggested to vary because of differential lignification, resulting in variations in hydrophobicity and, thus, in water repulsion [18]. This suggestion is consistent with the observation that the tendrils of various Cucurbitaceae species and the extracted lignified tissue ribbons therefrom coil further upon drying [18], [20].

The described free coiling movement concomitantly causes an axial shortening of the tendril. The tensional force generated in this process reduces the distance between the tendril’s insertion point at the climber’s stem and its anchorage point to the support, thus pulling and lashing the whole plant closely to its support structure (e.g., [7]) (fig. 1 b). The force generated by the free coiling movement, denoted as the coiling force in the following, was measured for tendrils of *Passiflora* and *Cucurbita* species many years ago by MacDougal [7], [10]; he used various dynamometer setups and found that the coiling force exceeded the plant stem’s weight and decreased in a wilted plant. More recently, Eberle et al. [23] employed a pendulum setup to measure the coiling force generated by *Luffa cylindrica*. Neither of these setups, however, included the continuous measurement of the force development. Continuous force-time curves for the coiling force generated by *Luffa cylindrica* have been reported by Liou et al. [24], but their cantilever wire setup did not discriminate the coiling force from the force exerted by the plant weight.

In this study, we present a custom-built setup that allows, for the first time, in situ measurements of tendril coiling forces over time, isolated from plant weight and in exact axial alignment. We have studied tendrils of *Passiflora caerulea* as a representative of the passion flower genus. Like the vast majority of the more than 500 passion flower species, *P. caerulea* is a vigorous climber that uses axillary unbranched tendrils for attachment via contact coiling [25], [26]. It is native to Argentina, Brazil, and Paraguay [25]. To complement the coiling force measurements and to gain a deeper understanding of the mechanism involved in force generation, we have analyzed tendril anatomy at various stages in ontogeny. Inspired by MacDougal’s [10] observation that the coiling force decreased in wilting plants and by Gerbode et al.’s [18] findings that both the entire cucumber tendril and the isolated fiber ribbons therefrom continue coiling even under drying, we have analyzed the dependence of the coiling force on the hydration status of *Passiflora caerulea* plants. We have therefore complemented our in situ measurements with dehydration-rehydration experiments on both entire tendrils and isolated lignified tissues.

In summary, this study aims to answer the following questions:

1. Do *P. caerulea* tendrils develop sufficient force to support and move the plant stem?
2. Is this force sensitive to fluctuations in water availability?
3. How do the different tissues contribute to the tendril coiling?

## Results

### Coiling force measurements

The coiling force of four *P. caerulea* tendrils was measured. From these, three measurements served primarily for studies of the maximum force produced and one measurement additionally served for investigating the influence of the plant’s hydration status on the coiling force. For this measurement, soil humidity was tracked continuously during tendril coiling, the coiled tendril was cut off from the plant and left to dry within the measurement setup and the coiling motion during the measurement was quantified (fig. 2).

**Fig. 2:**
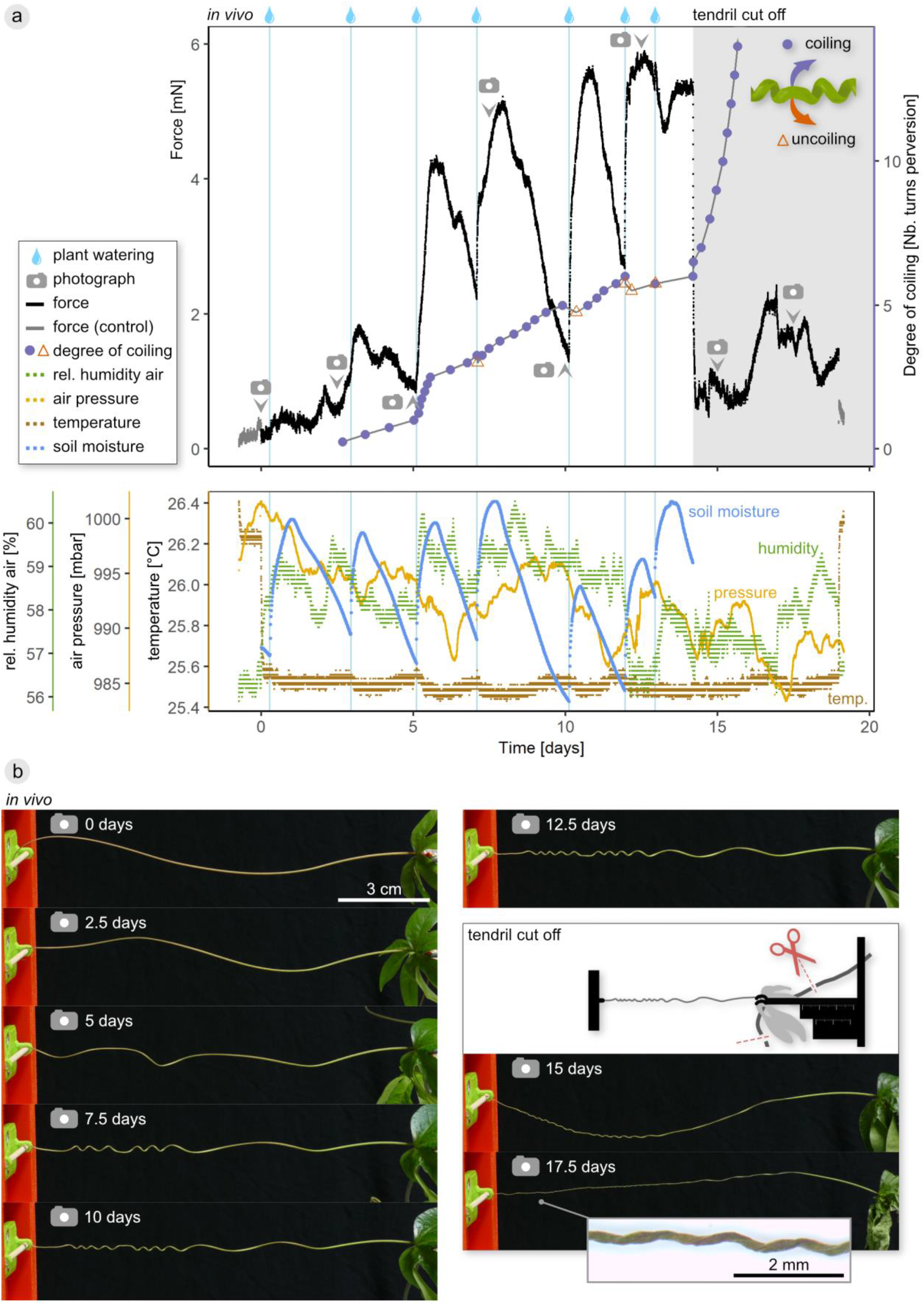
Tensile force generated during free coiling of a *P. caerulea* tendril. Force and environmental conditions and the degree of coiling or uncoiling are plotted over time **(a)**. Drop icons indicate the watering of the plant pot. After 14.2 days, the plant stem was cut apically and basally from the tendril insertion point, leaving the tendril fixed in an unaltered position in the experimental setup during drying. Photograph icons indicate video stills in **(b)**, inset shows magnified view of dried tendril. See electronic supplement video 2 for a synchronized video of force data and time-lapse video. Control indicates force before tendril was put in setup and after it was removed.

The coiling force in the four tendrils reached maximum total values of 6 mN, 11 mN, 49 mN, and 140 mN, respectively. The force initially remained low but increased when the tendrils started to coil (fig. 2, supplement fig. S1). Whereas in the two tendrils generating the highest maximum forces, the force remained high with only relatively small “jumps” upwards followed by a slower de-or increasing force, the force generated by the other two tendrils was subject to stronger fluctuations (fig. 2, supplement fig. S1). In general, the force-time curves were independent of fluctuations in temperature, relative humidity, or air pressure in the phytochamber.

The first sharp increase in force was correlated with an increased coiling movement expressed by a faster rotation of the perversion (fig. 2). Over time, the entire tendril axis coiled and formed one or more perversions, except for a basal segment that remained straight. Increases in force that occurred once the tendril had slightly coiled, i.e., established a connection under tension to the sensor, correlated well with the watering of the plant (fig. 2, see supplement fig. S1 for the other three measured tendrils). An increase in force was typically noticeable within less than 10 minutes after watering.

Once the tendril was coiled helically, both the increase and decrease in force was clearly correlated with the variation in soil moisture (fig. 2). Moreover, variations in the coiling motion coincided with watering: at the third watering point (day 5.1), the coiling speed increased sharply, whereas at the subsequent watering points (day 7.1, 10.1, 12.0, 12.9), the perversion slightly uncoiled (less than half a turn). At watering point five (day 10.1), the tendril had visibly drooped, and in parallel to the uncoiling of the perversion, it tightened again, i.e., the drooping tendril axis straightened. When the plant stem was cut apically and basally from the tendril insertion point at day 14.2, leaving the tendril fixed in an unaltered position in the experimental setup to dry off, the force started sharply to decrease within minutes. The coiled part of the tendril drooped, the coil shrank in diameter, and the number of coils increased quickly, causing the tendril axis to be lifted back upwards.

When a segment of the dehydrated tendril was submerged in tap water, it rehydrated and had an appearance similar to that before the tendril was cut from the plant stem (supplement fig. S2).

### Tendril anatomy

Within the coiled part of the tendrils, vascular bundles are arranged circularly or elliptically around the pith. The vascular cylinder is surrounded by the cortex, comprising of parenchyma and collenchyma, and the epidermis. Within the vascular bundles, primary vessel elements have lignified thickened cell walls that stain light blue with toluidine blue (TB) and red with phloroglucinol-hydrochloric acid (P-HCl) (fig. 3 a and b).

**Fig. 3:**
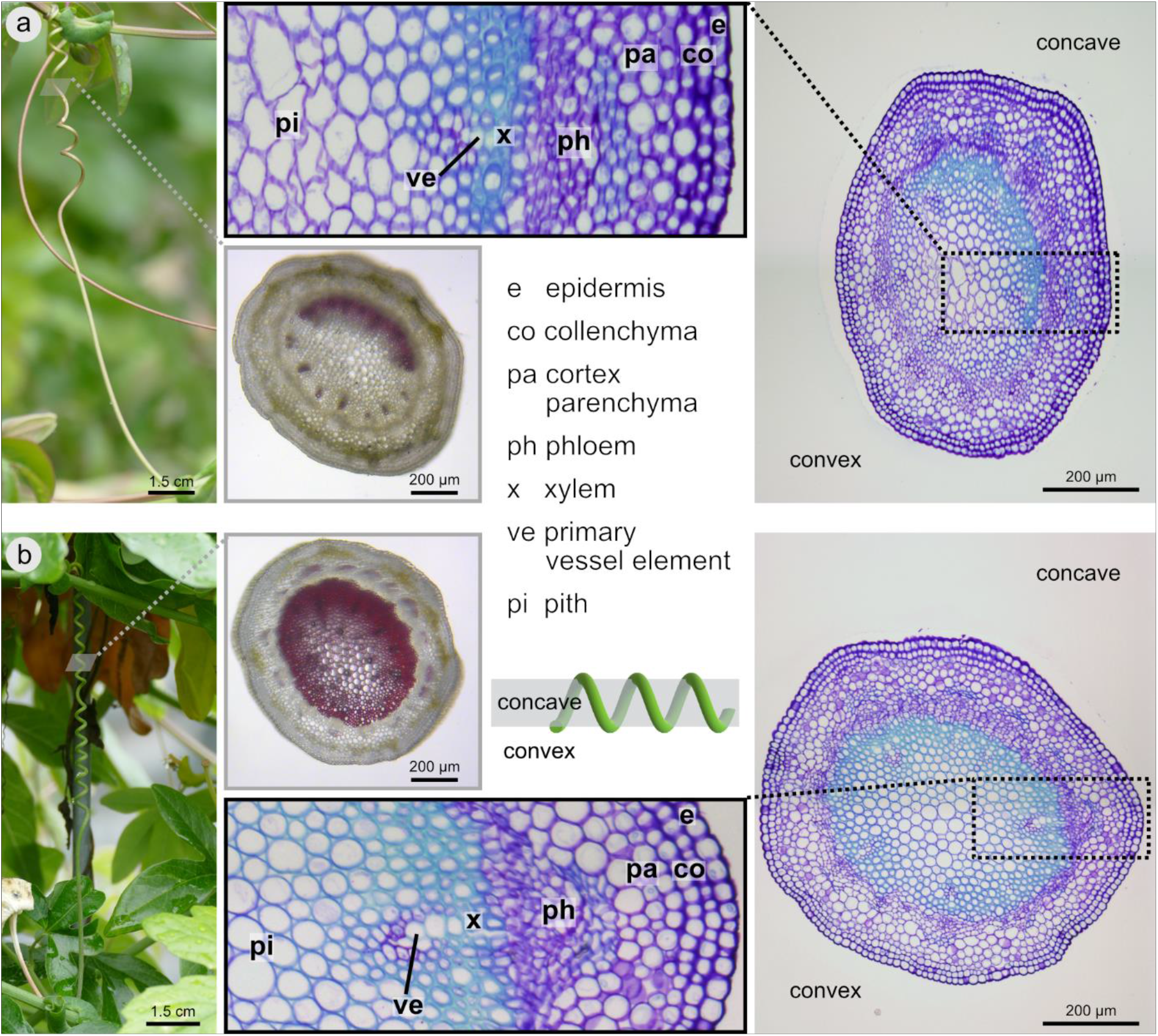
Tendril anatomy. Cross-sections through an immature tendril **(a)** showing red-(P-HCl) or light-blue-stained (TB) cells (evidence for lignification) in between the primary vessel elements on one side, which represents the concave side of the coiled tendril. In the mature tendril **(b)**, nearly the entire pith and xylem stained red (P-HCl) or light blue (TB). However, the concave side is still more intensely stained suggesting more intense lignification. Gray insets: Free-hand sections, stained with phloroglucinol-hydrochloric acid (P-HCl). Right side: thin sections of embedded material stained with toluidine blue (TB), with magnified black dotted insets (middle).

Based on the anatomical organization and differentiation, we distinguish between two ontogenetic stages: immature and mature tendrils. The former is characterized by thick-walled lignified cells, which stained light blue (TB) and red (P-HCl) (fig. 3 a), lying in between the primary vessel elements at one side of the tendril cross-section. This side corresponds to the concave inner side of the tendril coil. The pith parenchyma in the center and on the opposing outer side of the cross-section stained purple (TB) or remains unstained (P-HCl), indicating the lack of or limited lignification of the cell walls, and the cells are thin-walled. When treated with the cell wall degrading enzyme Driselase to remove parenchymatous tissues, a tissue with a crescent shaped cross section was isolated, representing a fiber ribbon (fig. 4 b1). Mature tendrils are characterized by a central cylinder comprising lignified xylem and pith, which stained completely red (P-HCl) and almost completely light blue (TB) (fig. 3 b). This central cylinder was isolated by treatment with Driselase (fig. 4 d1). The term “central cylinder” is used here as a descriptive term, rather than in the strict botanical sense.

**Fig. 4:**
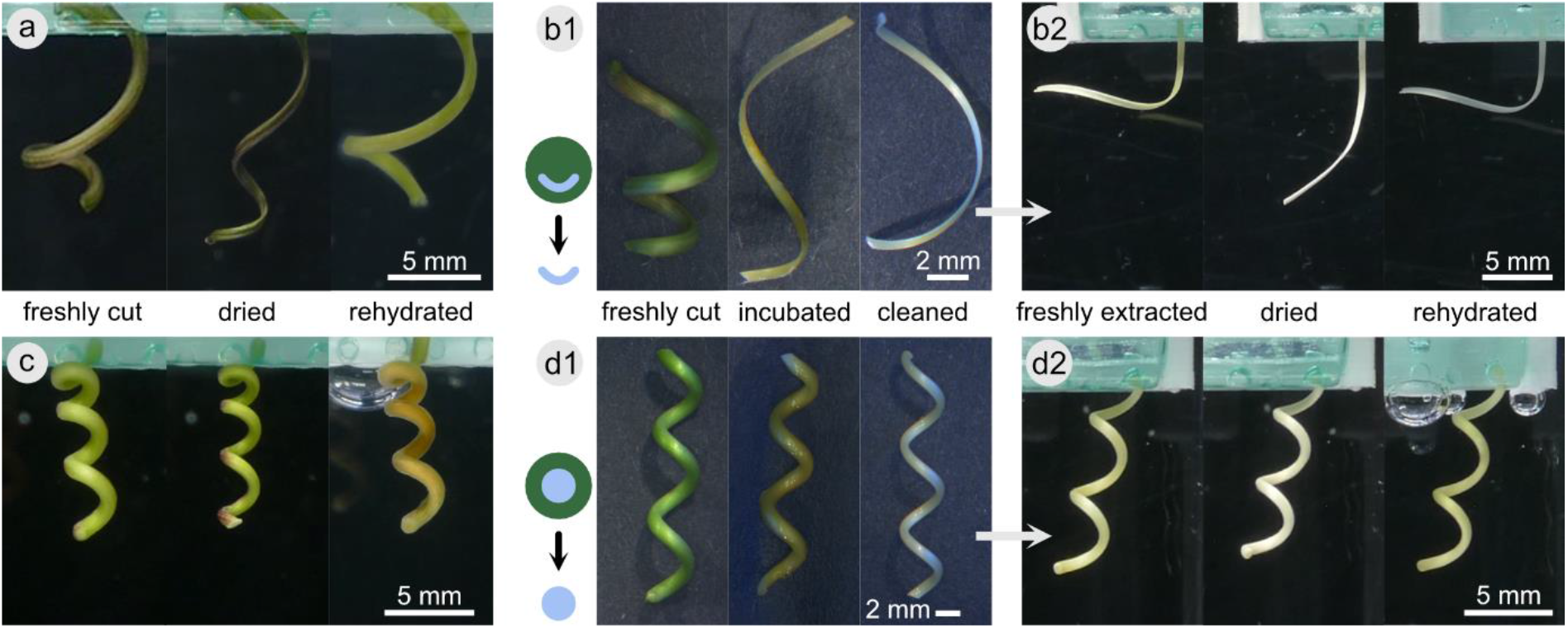
Tendril shape variations under de- and rehydration and extraction of lignified tissues. The effect of drying and rehydration on the shape of a coiled tendril segment is demonstrated for an immature tendril **(a)** and a mature tendril **(c)** and for extracted lignified tissues from an immature **(b)** and a mature tendril **(d)**. Stereomicroscopic pictures in the center (b1, d1) show a freshly cut tendril segment that was then incubated with the enzyme Driselase to digest parenchymatic tissues and cleaned to remove loose tissue remains. Images show the extracted samples or freshly cut samples in air, dried for two days in air, and rehydrated for two days submerged in water (three images in a, c, b2, d2, respectively). See also time-lapse videos of dehydration and rehydration in supplement video 3.

In the immature state, a large part of the tendril tissues collapsed when the tendril dehydrated. In comparison with the cross-section of a fresh (fully turgescent) immature tendril (fig. 3 a), the cross-section of a tendril that had been left to dry out in the coiling force setup (fig. 2 b) is distorted, and the exact tissue arrangement is hard to detect (supplement fig. S3). Only a small part of the cross-section stained light blue (TB), and the cross-section has a central cavity (fig. S3 a-c). In the basal straight part of the tendril, only the epidermis, cortex, and phloem had however collapsed, and the cross-section exhibits an intact central cylinder formed by a ring of mostly light-blue-stained (TB) and (in the outer part of the ring) thick-walled cells around a central cavity (fig. S3 d).

### De- and rehydration behavior

When an immature tendril segment was left to dry out, it underwent a complex shape-changing pattern and eventually coiled slowly, while its cross-section flattened (fig. 4 a and supplement video 3). Rehydrated, the tendril resumed a shape close to that of the initial one. When the lignified fiber ribbon from such an immature tendril was isolated by Driselase treatment, its coiled shape loosened, and only a smaller degree of curvature was maintained (fig. 4 b1). Left to dry, the isolated ribbon bent open. When rehydrated, it resumed its initial slightly curved shape (fig. 4 b2).

When a mature tendril was left to dry, it shrank in diameter and coiled by an additional small extent (fig. 4 c and supplement video 3). Upon rehydration, it resumed its initial shape. When the lignified central cylinder from such a tendril was isolated by Driselase treatment, it remained unaltered in its coiled shape (fig. 4 d1). Upon drying, the isolated cylinder contracted and coiled by an additional small extent. When rehydrated, it resumed its initial shape (fig. 4 d2).

### Fresh weight

The plant stems with leaves had a mass of 0.7 ± 0.2 g per node (mean ± standard deviation), i.e., per tendril, equivalent to 6.9 mN. The flower’s mean mass was 3.2 ± 0.5 g, equivalent to 31.4 mN.

## Discussion

We found that on coiling, the tendrils of *P. caerulea* generate tensional forces ranging from 6 to 140 mN. In nature, the tension caused by coiling transcribes to a shortening of the tendril by as much as one-third of its straight length for *P. caerulea* [7]. This is in good agreement with measurements performed by MacDougal [7], [10], who used a so-called Vöchting’s dynamometer and measured tensional forces between 30 and 100 mN for tendrils of *P. caerulea* and *P. pfordti*. Liou et al. [24], using a setup in which the shortening tendril bent a wire from which force was then extrapolated, measured tensile forces of 170 mN for *Luffa cylindrica* tendrils, whereas Eberle et al. [23] measured only 3.4 ± 0.2 mN of tensional force for the same species, by using a pendulum apparatus. The latter eliminated the influence of the plant’s own weight on the measurements by restraining the plant stem. Our study is the first that allows continuous measurement of the tensional force that coiling tendrils are able to generate when both ends are fixed. As the plant stem produces about 7 ± 2 mN of weight force per node and thus per tendril, the generated tensional force indeed enables the tendrils to move the plant stem against gravity and thus to lash the plant to its support.

Tendrils coil because of their intrinsic curvature (e.g., [8]), which is generated by a length difference between the tendril’s sides with the convex side being longer than the concave side. Since contractile G-fibers are common in climbing plant tendrils ([17], [12], own unpublished results fig. 5), we can safely assume that contraction is involved in the build-up of the length difference. The most straightforward explanation for the occurrence of the observed tendril bending in *P. caerulea* is thus the existence of a bilayer composed of an actuating contractile layer and a second antagonizing non-(or less) contractile resistance layer. The coiling and formation of perversions by such a bilayer can be visualized, for example, by stretching silicone rubber, adding a second layer of relaxed silicone, and releasing the thus fabricated bilayer strips (e.g., [18]).

**Fig. 5:**
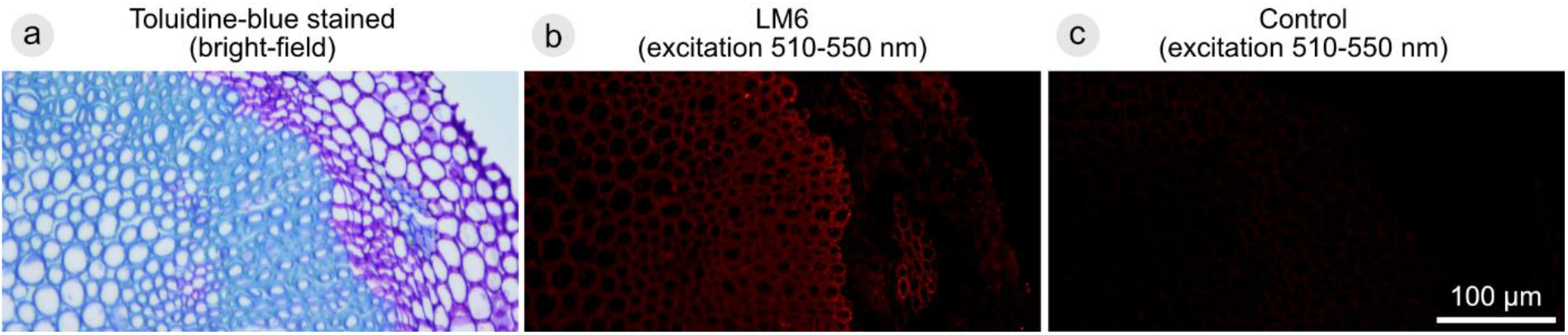
Immunolocalization of a G-layer polysaccharide in fiber cells of *P. caerulea* tendrils. Detail of cross-sections through a mature tendril stained using toluidine blue **(a)** or via indirect immunolabeling with the antibody LM6 **(b)**. The red fluorescent signal indicates the localization of LM6, which targets RG-I (rhamnogalacturonan-I). RG-I is a pectic polysaccharide found in plant primary cell walls and in the G-layer of fiber cells [17], [31], [32]. The strong signal in the tendril central cylinder indicates the presence of G-fibers. Control in **(c)** without addition of antibody LM6.

The tissue organization in mature *P. caerulea* tendrils, however, is not bilateral, but rather centri-symmetric, comparable to redvine tendrils (*Brunnichia ovata*), wild grape tendrils (*Vitis rotundifolia*), or greenbrier tendrils (*Smilax rotundifolia*), for which G-fibers have been reported in a hollow cylinder, or to tendrils of the Virginia creeper (*Parthenocissus quinquefolium*), in which they are positioned in the central region of the tendril [12]. A contraction of a centri-symmetric tissue, however, will not lead to curvature and coiling. As a consequence, the anatomy of the tendrils at the time of the formation of the helical coils cannot have been centri-symmetrical but becomes centri-symmetrical only later in the maturation process. Indeed, our results show that, in *P. caerulea* tendrils, a bilateral tissue organization develops as a transient structure during a young ontogenetic phase. In young straight tendrils, the primary vessel elements are the only lignified structures [29], but during coiling, the xylem starts to lignify unilaterally on the concave side of the tendril. This concurs with observations by MacDougal [30] who described, more than a century ago, the unilateral development of wood at this side. In the basal tendril region, which does not coil at all, the xylem is arranged symmetrically, even in the immature tendril state (fig. S3 d, cf. also [30]). The obvious bilateral tissue organization in the coiling part of the young developing tendrils shows greater similarity to cucurbit tendrils in which G-fibers occur as a ribbon on the concave side of the tendrils [12].

The lignified fiber ribbon is specific for the coiling part of the tendril of *P. caerulea* and lies at a position similar to the G-fiber ribbon described in cucurbit tendrils. Based on this finding, we assume that, in *P. caerulea*, the maturation of the fiber ribbon itself drives the coiling of the tendril, most likely by providing contractile properties via a G-layer in the fiber cells. So far, G-fibers have been reported in *Passiflora* sp. tendrils ([18], supplements) and in stems of *P. caerulea* ([19], dataset S1), albeit in an extra-xylary position. With regard to *P. caerulea* tendrils, we have first histological evidence for the existence of a G-layer in the fiber cells (unpublished data) (fig. 5). However, this needs to be confirmed by more targeted histological analyses for G-layer detection, including immunolabeling or G-fiber identification based on morphology and the differential staining of the cell wall layers [19]; this was beyond the scope of this study and is the subject of ongoing research.

The exact mechanism by which G-fibers generate tension remains under discussion (e.g., [14]). Tension wood, i.e., wood that has a G-layer, is known strongly to shrink in a longitudinal direction when dried (summarized in [14]). Consistently, dehydration of the hydrophilic G-layer has been suggested as a driving factor for G-fiber contraction [31], although this model has been rejected by others [14]. Independent of the exact mechanism causing the active G-fiber contraction involved in tendril coiling, we assume that contraction can also be induced passively by drying of the G-fibers (cf. [18], [16]). An extracted fiber ribbon from a *P. caerulea* tendril, however, does not coil when dried, indicating that it represents only one individual layer (the actuating layer) of the bilayer necessary for tendril coiling. This is unlike the situation in cucumber tendrils in which the G-fiber ribbon alone coils when dried, as it constitutes an asymmetrically contracting bilayer that imposes its coiled shape on the surrounding soft tissue [18].

Overall, our data strongly suggest that the tendrils of *P. caerulea* actively coil because of the contraction of an eccentrically positioned G-fiber ribbon, antagonized by the surrounding compression-resistant parenchyma, which acts as a resistance layer in the turgid state. The thick parenchymatous tissue layer on the convex side of the fiber ribbon resists contraction more strongly than the thinner layer on the concave side, and the tendril thus bends and coils. The fiber ribbon develops during the phase of active movement in the immature tendrils, namely a phase that represents a transitional phase in tendril ontogeny, and eventually leads to the lashing of the plant to its support and the building of a spring-like shape.

The tension generated by tendril coiling correlates strongly with the watering of the plant, which we assume is translated (with a certain delay) into turgor pressure within the cells of the tendril. During coiling, the tendrils remain in an immature state and consist mostly of thin-walled parenchyma cells. Parenchymatous tissues mechanically act as hydrostats, whose cell walls are placed under tension by the pressure generated within the walls [33]. The thus “inflated”, fully turgescent, parenchyma cells are compression-resistant and resist external deformation, whereas at lower internal pressures, the tissue becomes less stiff and ultimately flaccid. When the turgor pressure and thus the (compressive) stiffness of the parenchyma tissue decreases, the stiffness of the entire spring-like tendril also decreases, and, as a consequence, the helical coil relaxes. The tendril-spring slackens, and the measured tensional forces decrease. When the turgor pressure of the parenchyma cells increases after the watering of the plant, however, the spring is placed under tension and raises the tendril axis minimally but rapidly, a movement that can even slightly uncoil the tendril as observed in our experiments. This reaction to watering occurs rapidly, within several minutes, and indicates efficient water transport from the soil to the tendril. This is in agreement with evidence that lianas are highly efficient in being able to conduct water [34], [35], [36]. In addition to the parenchyma, the tendrils contain the lignified fibers that were discussed above. Although lignification stiffens the cell walls [33], the thin ribbon-shaped lignified fiber strand found in young immature tendrils can relatively easily twist and bend when the G-layers of the fibers contract and thereby causing coiling in interaction with the parenchymatous resistance layer. Even though these measurements show impressively that the tension in the tendrils during the coiling phase depends directly on the water balance of the plant itself, we can assume that this has no vital influence on the attachment of the stem or the plant as a whole. In older, fully developed, ontogenetic stages, the tendrils, that are now almost fully lignified, die off and thus become entirely water independent, while still contributing to the overall attachment of the plant stem.

When parenchyma cells of an immature tendril become flaccid because of complete turgor loss, the generated tension is lost. During drying, the parenchyma becomes less stiff because of turgor loss, but also collapses causing the tendril diameter to shrink and the cross-sectional geometry to distort. Moreover, the tendril coils further, so that overall, the spring’s pitch and radius change dramatically. The resulting “spring” (fig. 2 b) probably not only loses its intrinsic tension, but also its energy dissipating capacity.

This dehydration-induced coiling is a passive and reversible motion based on the tissues’ hygroscopic properties. Hygroscopic (i.e., humidity-driven) movements can be found across the Plant Kingdom, often being associated with the dispersal of seeds and spores. This includes, for example, the opening and closing of pinecones [37], [38] and silver thistle bracts [39], the dispersal of spores in mosses [40], the penetration of seeds into soil [41], and the opening of seedpods [42], [43]. Hygroscopic movements rely on dead cells, whose swelling/deswelling behavior is dictated by cellulose microfibril orientation in their cell walls [44]. When layers are combined that have variable cell wall architecture or cell orientation, and thus variable preferred directions of contraction, the dehydration of a tissue can lead to bending, twisting, or coiling [45]. Even dehydration of a specialized single homogeneous layer can induce coiling, as has been shown for *Erodium* awns [46]. As all these movements merely rely on humidity, they occur passively, i.e., without requiring metabolic energy. The free coiling movement in *P. caerulea* tendrils, however, which changes a straight tendril into a coiled and tensioned tendril spring, is an active (i.e., metabolically costly) movement that relies on the maturation of G-fibers and on live hydrostatic parenchyma cells. Nevertheless, the passive additional coiling upon drying suggests that differential dehydration behavior also plays a role in active coiling.

Unlike *Bauhinia* seedpod valves or *Erodium* awns, which coil only in one handedness, coiling tendrils, which are fixed at both ends, form helices of opposite handedness and one or several perversion points within one organ. Thus, the helix formation is unlikely to result directly from the twisting or coiling of individual cells or tissues, as this would require differential handedness in the cell walls, cells or tissues along the tendril. Instead, as discussed above, the coiling process is driven by contraction along the entire organ, which induces curvature and thereby coiling. The precise mechanism behind the hygroscopic coiling motion remains to be determined. A better understanding of this mechanism would be valuable not only from a purely botanical viewpoint, but also from a material science perspective in order to explore the tendril as a hydration-actuated shape-memory structure.

During ontogeny, lignification proceeds across the entire central cylinder. In contrast to the thin ribbon, the resulting lignified cylinder now fully retains its coiled shape when isolated, i.e., the spring shape in fully developed mature tendrils is structurally “frozen”. Lignin itself is not particularly resistant to tension stress, but the presence of lignin reduces water infiltration and the water content of the cell walls [33]. Since dry cellulose is considerably stiffer than wet cellulose, lignification stiffens the cell wall and stabilizes its mechanical properties against fluctuations in water content [33]. As a consequence, the tension generated by the coiling tendril becomes increasingly independent of the water status of the plant as lignification proceeds during maturation. Furthermore, with the increasing stiffness of tissues comprising the tendril, the stiffness of the tendril-spring as a whole also increases. A stiffer spring can carry heavier loads and provides better energy dissipation, since it deflects less under a given load. This is important as the plant weight increases over time because of leaf growth, the emergence of flowers, and the production of fruits.

In contrast to the hydrostatic, unlignified, thin-walled parenchyma cells, the thick-walled lignified cells do not collapse when dehydrated, so that in the mature state, the spring-shape of the tendril barely changes under water stress. As is known from other passion flower species, the tendrils, in a final step during ontogeny, dehydrate and completely dry out in the natural process of senescence, while still maintaining a high degree of functionality [22]. This can be interpreted as an “energy-saving mechanism” minimizing the long-term metabolic energy costs for the plant. Observations of adult plants in the Botanic Garden Freiburg show that in *P. caerulea*, subsequent to the mature state investigated here, the tendrils become senescent while maintaining mechanical functionality. Lignification of the full central cylinder might indeed be the key to enable this dehydration, while avoiding large changes in spring-shape and, thus, significant losses in mechanical functionality.

In brief, the tendril springs in *P. caerulea* form in two phases (fig. 6). First, during an active movement phase after the tendril made contact with a host, it hoists the plant stem toward the host in an upward direction, and forms its spring-like shape. This phase then transitions into a stabilization phase in which the entire central cylinder lignifies. This two-step process has inspired the development of artificial tendril and attachment devices (e.g., [47], [29]).

**Fig. 6:**
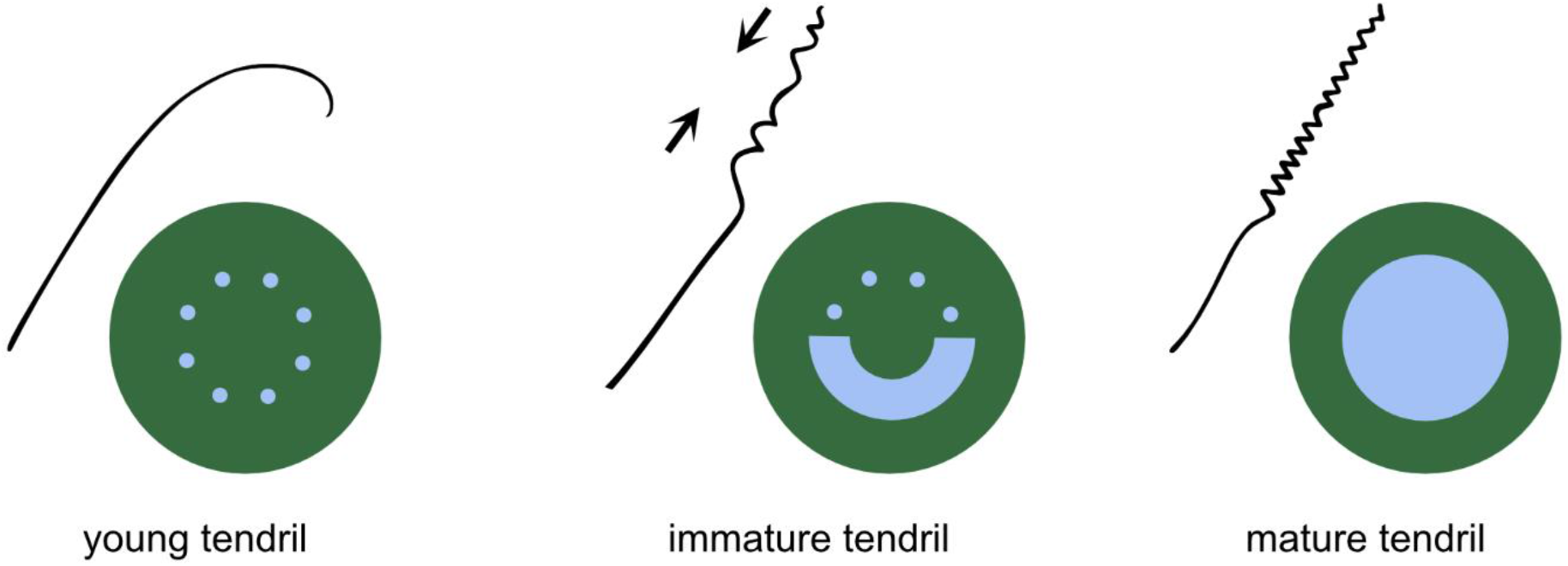
Model for two-phasic spring formation in *P. caerulea* tendrils. After attachment of the tendril tip, the immature tendril first undergoes an active phase of motion, during which it coils along its axis and thus lashes the plant to its support. The coiling is an active motion, probably driven by the contraction of G-fibers, observed as a lignified ribbon on the concave side of the tendrils. In a second phase, the spring-shape resulting from the coiling is stabilized and further stiffened by lignification of the entire central cylinder. The representations simplify the distribution of lignified (light blue) and parenchymatous (green) tissues.

## Conclusion

We have shown that the coiling tendrils of *P. caerulea* generate forces large enough to move a plant stem and to carry its load. Our data suggest that the tendrils coil by the active contraction of an eccentrically positioned G-fiber ribbon functioning together with the surrounding antagonizing parenchymatous tissues acting as resistance layer in a turgid state. Rather than being a persistent shape, the ribbon shape of the lignified fiber strand merely represents a transitional ontogenetic phase. In this immature state, the generated tension strongly varies with the water status of the plant, i.e., its turgor pressure, since the tendril spring is predominantly parenchymatous, and the hydrostatic stiffness of parenchyma depends on its turgor pressure. In addition to active coiling, the tendril in this state also coils significantly when undergoing drying, i.e., in a passive hygroscopic way. However, it loses its mechanical functionality when drying, as the initial taut spring shape is lost, as is the generated tension and the spring properties.

In a second ontogenetic phase, which can be regarded as a stabilization phase, the lignification progresses beyond the fiber ribbon, so that the shape of the lignified tissue changes from a relatively thin ribbon to a full cylinder. This renders the shape of the spring independent of the existence of a hydrated parenchyma, probably entailing several advantages: (i) the generated force becomes more independent of the turgor pressure, (ii) the spring stiffens, i.e., it can bear greater loads, and (iii) the spring dehydrates with a much smaller shape change, so that it can retain a higher amount of mechanical functionality when becoming senescent. With their ontogenetic multi-functionality and hydration-actuated shape changes, the coiling tendrils thus provide several intriguing features that should be explored for biomimetic transfer extending beyond the coiling motion itself.

## Methods

### Plant cultivation

For measurements of the coiling force a potted plant of *Passiflora caerulea* was cultivated in a phytochamber, under controlled temperature and humidity conditions (25.7±0.2 °C, 57±1%; mean and standard deviation for conditions logged in the vicinity of the plant pot during coiling force measurements, data logger EL-SIE-6+, Lascar electronics, Whiteparish, United Kingdom; accuracy ±0.2 °C, ±0.1%). The plant was grown under constant illumination during the entire experimental time, i.e., without any dark periods (UGR19 Highbay RKU-200W-NW LEDs, LeuchTek GmbH, Ahrensburg, Germany), and the light level at the plant was between 120-140 μmol m^−2^ s^−1^ PPFD (light meter LI-250, LI-COR Biosciences GmbH, Lincoln, USA). Mealybugs were occasionally spotted on the plants but were immediately brushed off using a paintbrush. Shoots for the measurement of their fresh weight were collected from potted *P. caerulea* plants grown in the greenhouse of the Botanic Garden of the University of Freiburg, and flowers from the potted plant cultivated in the phytochamber. For histology and dehydration-rehydration experiments, tendrils were collected from *P. caerulea* plants cultivated in either the greenhouses or outdoor areas of the Botanic Garden.

### Coiling force measurement

The coiling force of four *P. caerulea* tendrils was measured using a custom-build setup (fig. 7). Whereas three measurements served primarily for studies of the maximum force produced, one tendril was analyzed with the aim of additionally investigating the influence of the plant’s hydration status on the coiling force. During this measurement, soil humidity was tracked continuously during tendril coiling, and the coiled tendril was cut off from the plant and left to dry within the setup.

**Fig. 7:**
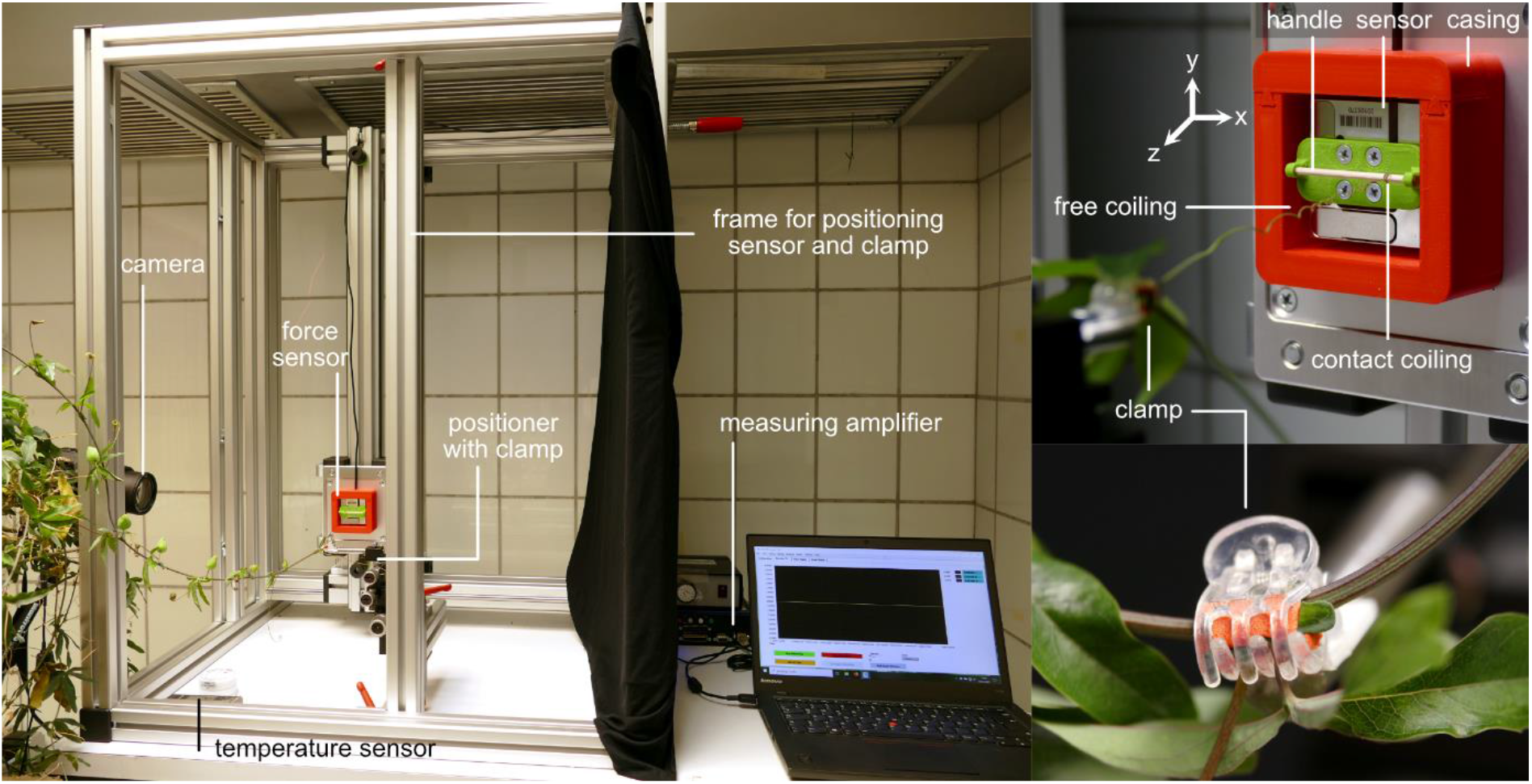
Experimental setup. To measure the force generated by the free coiling of *P. caerulea* tendrils, a custom-built measuring setup consisting of a 3-axes-force sensor was used, which was equipped with a handle, around which the tendril apex coiled (contact coiling), and a positioner with a clamp, which retained the stem at the tendril base.

The setup consisted of a 3-axis-force sensor (K3D40 ±2N, accuracy class 0.5%), in combination with a measuring amplifier (GSV-8DS SubD44HD, both ME-Meßsysteme GmbH, Henningsdorf, Germany) and the manufacturer’s software (GSVmulti version 1.46.1 2020) (fig. 7). The sensor was attached to a movable mount within a sturdy aluminum frame, allowing the free positioning of the sensor within a 2D-plane. On the opposite side of the frame, a micro-positioner was positioned equipped with a 1 cm hairgrip clamp padded with a piece of craft foam sheet, to immobilize the shoot and thus to prevent it from tensioning the tendril by its own weight. A custom-made 3D-printed holder (ABS) with a small handle consisting of a bamboo toothpick (Wenco-Service Marketing GmbH & Co. KG, Essen, Germany) was mounted on the live end of the sensor by using four cylindrical head screws. The handle was 2 mm in diameter and 37 mm in length, and the distance between the handle’s central axis and the back of the holder, i.e., the sensor surface, was 6.5 mm. An uncoiled tendril with a slightly bent tip was carefully and loosely placed on the handle. The stem was clamped at the tendril’s base. Both the apex and the leaf at the clamped node developed normally, i.e., clamping did not disturb the development of the plant. The tendril was continuously photographed at a frequency of 1 image per minute (Lumix DMC-FZ1000, Panasonic, Kadoma, Japan), and the individual images were combined to produce a time-lapse video. Temperature, relative humidity, and air pressure were logged at 1/60 Hz (data logger EL-SIE-6+, Lascar electronics, Whiteparish, United Kingdom; accuracy ±0.2 °C, ±0.1%, ±1 mbar). All potential disturbances of the measurement (by people entering the chamber to attend to the experiments or to water the plant pots) were carefully noted. The force along each of the x-, y- and z-axis was measured with a sampling frequency of 1 Hz, which was reduced to 1/60 Hz for subsequent analysis of the data. From the three force channels, the total force parallel to the coiled tendrils length axis was calculated as 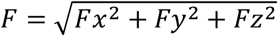.

For the measurement targeting the hydration effects on the coiling force, the soil humidity in the plant pot was continuously tracked (one measurement every 15 minutes; sensor TMS-4, TOMST s.r.o., Prague, Czech Republic, [27]; the data regarding the raw moisture count are reported). To quantify the degree of coiling, the rotational motion of the perversion visible in the time-lapse video (supplement video 2) was analyzed, and the number of turns that it performed was counted in 0.25 steps. Any reversal in the turning direction of the perversion was documented (uncoiling), and the turns were counted, rounded to 0.25 turns. After 14.2 days, each plant stem was cut apically and basally to the clamped section leaving the coiled and still clamped tendril to dry out in its original position and thus allowing the force measurement to be continued. With regard to the drying tendril, the degree of coiling was quantified as described above, except that the number of turns that the perversion performed was counted in larger steps. After the tendril had been left to dry in the setup for 4.8 days, it was removed from the setup for further analyses in the laboratory. A small segment from the coiled part of the dried tendril was rehydrated in tap water for 30 min and photographed (Lumix DMC-FZ1000, Panasonic, Kadoma, Japan). From the remaining dried tendril, three segments from the coiled part and one segment from the straight basal part were embedded and thin sectioned, as described in the following section.

### Histology

Based on preliminary experiments, we distinguished between immature and mature tendrils depending on the development of their anatomical features (cf. results section). From each developmental stage, two tendrils were collected for lignin-specific phloroglucinol-hydrochloric acid staining of fresh sections [28]. Parts of the same tendrils were also used for dehydration-rehydration experiments (see next section). An additional tendril of each developmental stage was collected for embedding in resin, thereby allowing a precise alignment of the coiled part of the tendril with respect to its length axis. All tendrils were sectioned within the coiled tendril part.

In the discussion, we mention initial results obtained from parallel, still unpublished, research, with a more in-depth focus on the anatomical development of *Passiflora* tendrils during coiling. More precisely, we report the immunolocalization of RG-I (rhamnogalacturonan-I) in the cross-sections of a fully coiled *P. caerulea* tendril, by using the primary antibody LM6 ([Anti-1,5-α-L-Arabinan] Antibody, Megazyme Ltd, Bray, Ireland) and a fluorescent marker (Alexa Fluor 568 goat anti-rat IgG (H+L), Thermo Fisher Scientific Inc., Waltham, United States). The supplement provides the full protocol.

For lignin detection, fresh sections were cut with a razor blade and were subsequently submerged briefly in a phloroglucinol-ethanol solution (wt 1%) and then briefly in 20% hydrochloric acid. For microscopy, sections were transferred to tap water on a slide and photographed (microscope Primostar with camera Axiocam ERc 5s, both Zeiss, Oberkochen, Germany).

For thin sectioning of embedded samples, tendril material was fixed in FAA, namely a mixture of 70% ethanol, 37% formaldehyde solution, and glacial acetic acid (at a ratio of 90:5:5) for 2-48 days. Samples were then dehydrated through a series of increasing isopropanol concentrations and embedded in resin (Technovit 7100, Kulzer GmbH, Hanau, Germany). Sections were obtained using a rotary microtome (CUT 5062, SLEE medical GmbH, Nieder-Olm, Germany), stained with toluidine blue (0.05% (wt)), and transferred to glass slides. For microscopy, the slides were sealed with Entellan (Merck, Darmstadt, Germany), and the sections were photographed (microscope BX61 with camera DP71, both Olympus, Tokio, Japan).

### Dehydration-rehydration experiments

To investigate the effect of the hydration status on the shape of the coiled tendril and on isolated lignified structures, two mature and immature tendrils were collected. A coiled segment was cut from the distal end of each tendril. Lignified tissues were isolated from one tendril per ontogenetic state. To this end, the segment was incubated with the enzyme Driselase for 72 hours at 22°C-37°C (based on [18]) (Sigma-Aldrich, Missouri, USA; 2% (wt) Driselase solved in phosphate-buffered saline). Samples were then sonicated for 20 min (ultrasonic cleaner Emmi H60, EMAG AG, Mörfelden-Walldorf, Germany), and any remaining parenchyma tissue material stuck to the lignified structures was gently removed using a brush. The isolated lignified structure was photographed (cf. fig. 4 b1 and d1) (stereomicroscope SZX9, Olympus, Tokio, Japan with camera Color View II, Soft Imaging System GmbH, Münster, Germany). Both the treated tendril segments and the fresh segments from each developmental stage were fixed at one end, air-dried for 48 hours, and subsequently submerged in tap water for another 48 hours. The drying and rehydration process was photographed (1 picture per minute, Lumix DMC-FZ1000, Panasonic, Kadoma, Japan).

### Measurement of fresh weight of plant material

To determine the fresh weight of the plant material, 37 plant stem segments of 1 m were harvested, including the leaves. The number of nodes in each segment was counted, and the segment was weighed, enabling the mean weight per node to be determined. Additionally, nine flowers with stalks were cut and weighed.

## Data availability statement

All data supporting the findings of this study are available within the paper and its supplementary information files, or are available from the corresponding author (Frederike Klimm) upon reasonable request.

## Acknowledgements

This work was supported by the European Union’s Horizon 2020 Research and Innovation Programme under Grant Agreement No 824074 within the framework of “GrowBot - Towards a new generation of plant-inspired growing artefacts”. TS and FK acknowledge additional funding by the Deutsche Forschungsgemeinschaft (DFG, German Research Foundation) under Germany’s Excellence Strategy-EXC-2193/1-390951807 and access to the shared laboratory facilities of the cluster.

The authors thank all at the technical workshop of the faculty Biology II/III for their support with the measurement setup, Michael Schmidt for the preparation of thin-sections, and Michelle Modert for immunolabeling.

## Author contributions

TS and MT: conceptualization and supervision; TS: funding acquisition; FK: measurements; FK and MT: data evaluation; all authors: discussion and interpretation of data; FK: writing the first draft; TS and MT: improving further versions. All authors gave final approval for publication.

## Materials & Correspondence

Correspondence and requests for materials should be addressed to Frederike Klimm.

## Supplementary data

- Protocol P1: Immunolocalization of RG-I in a cross-section of a tendril of *P. caerulea*
- Fig. S1: Tensile force generated during free coiling of *P. caerulea* tendrils
- Fig. S2: Rehydration of a dehydrated *P. caerulea* tendril segment
- Fig. S3: Tendril anatomy of a dehydrated *P. caerulea* tendril
- Video V1: Coiling *P. caerulea* tendril (complementing fig. 1)
- Video V2: Force generation in a coiling *P. caerulea* tendril (complementing fig. 2)
- Video V3: Dehydration-rehydration behavior of *P. caerulea* tendrils (complementing fig. 4)

