## Supplementary material for "Force generation in the coiling tendrils of *Passiflora caerulea*"

**Protocol P1: Immunolocalization of RG-I in a cross-section of a tendril of *P. caerulea***

A fully coiled tendril was cut to size and fixed overnight in FAA, dehydrated in a series of increasing acetone concentrations, embedded in Technovit 8100 (Heraeus Kulzer GmbH, Wehrheim, Germany), and sectioned on a rotary microtome. In order to localize the distribution of RG-I (rhamnogalacturonan-I) in the tendril cross-section, we used indirect immunolabeling (fig. P1). Sections were treated with a primary antibody targeted at the arabinan side chains of RG-I (LM6 [Anti-1,5-α-L-Arabinan] Antibody, Megazyme Ltd, Bray, Ireland). The location of the primary antibody was visualized using a secondary antibody with a fluorescent marker (Alexa Fluor 568 goat anti-rat IgG (H+L), Thermo Fisher Scientific Inc., Waltham, United States).

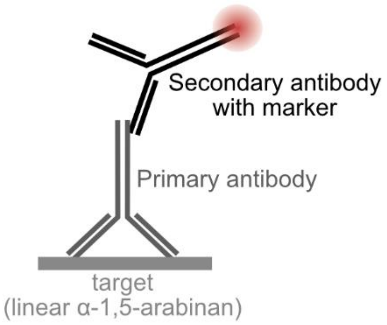


**Fig. P1: Principle of indirect immunolabeling**. Sections were incubated with a primary antibody targeted at the arabinan side chains of RG-I, which was then visualized using a secondary antibody with a fluorescent marker.

For immunolabeling, slides were processed as follows:

- Phosphate-buffered saline (PBS) (10 min)
- 0.3 M glycine in PBS (20 min)
- 3% bovine serum albumin (BSA) in PBS (30 min)
- Primary antibody, diluted 1:20 in 3% BSA in PBS (overnight at 4°C)
- Phosphate-buffered saline with Tween 20 (1:1000) (PBST) (5 min, 3x)
- Secondary antibody diluted 1:200 in 3% BSA in PBS (60 min)
- PBST (5 min, 3x)

As a control, slides were processed as follows:

- PBS (10 min)
- 0.3 M glycine in PBS (20 min)
- 3% BSA in PBS (30 min)
- Secondary antibody diluted 1:200 in 3% BSA in PBS (60 min)
- PBST (5 min, 3x)

Slides were treated with antifade medium (SlowFade Gold antifade reagent with DAPI, Thermo Fisher Scientific Inc., Waltham, United States), covered, sealed, and examined via an epifluorescence microscope (microscope BX61 with camera DP71; Olympus, Tokio, Japan). Antibody detection was carried out using an U-MWG2 mirror unit (excitation filter: 510-550 nm, dichroic mirror: 570 nm, barrier filter: 590 nm).

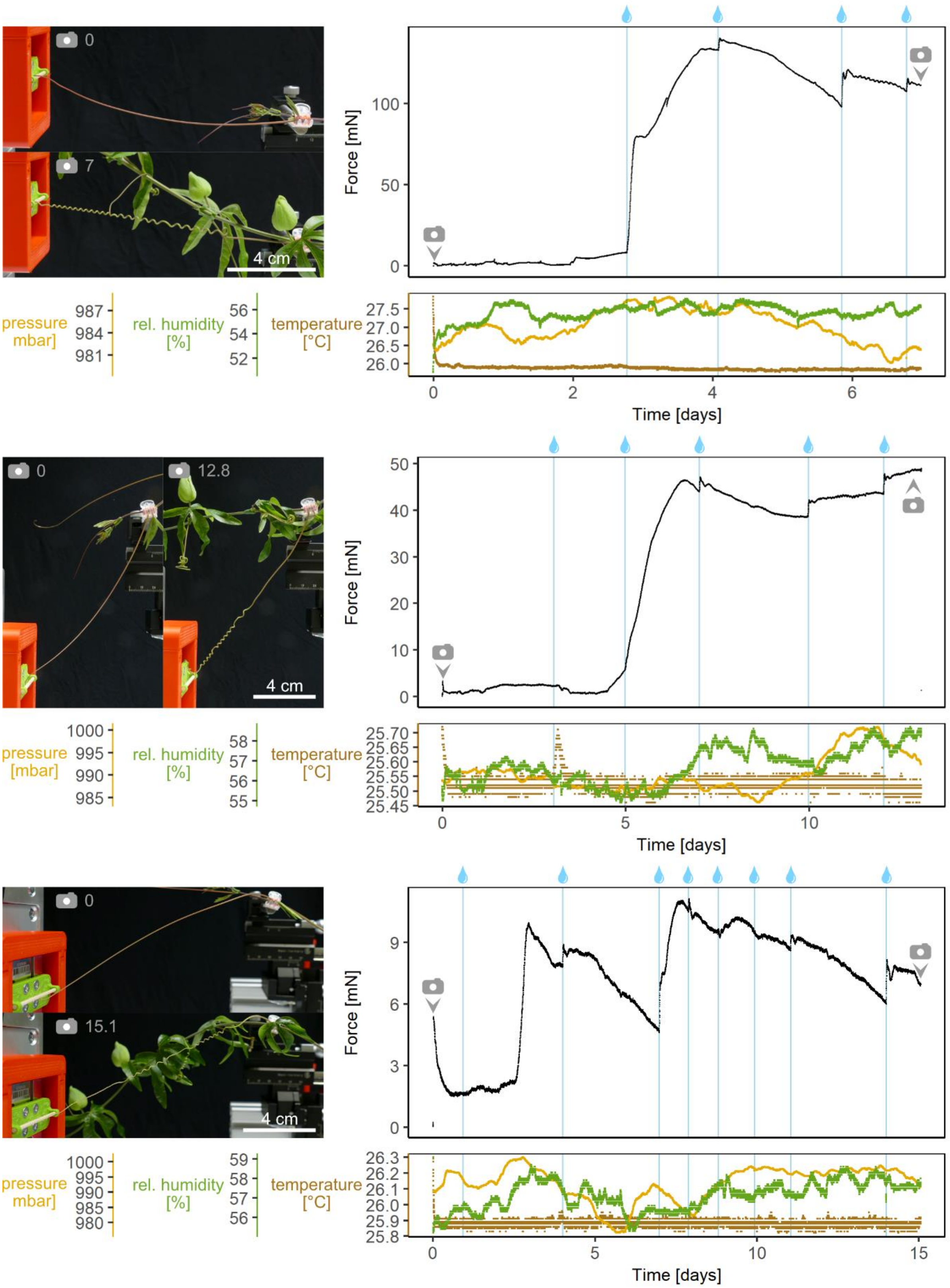


**Fig. S1: Tensile force generated during free coiling of *P. caerulea* tendrils**. Force and environmental conditions are plotted over time. Drop icons indicate the times at which the plants were watered, and camera icons indicate the time at which video stills were taken at the beginning and end of the measurements. For the third tendril, the initial load reflects the pre-tension caused by the clamping of the tendril base. Since the bent tendril tip opened up again before tightly grasping the handle, the force decreased before coiling started, and so the measurement is valid.

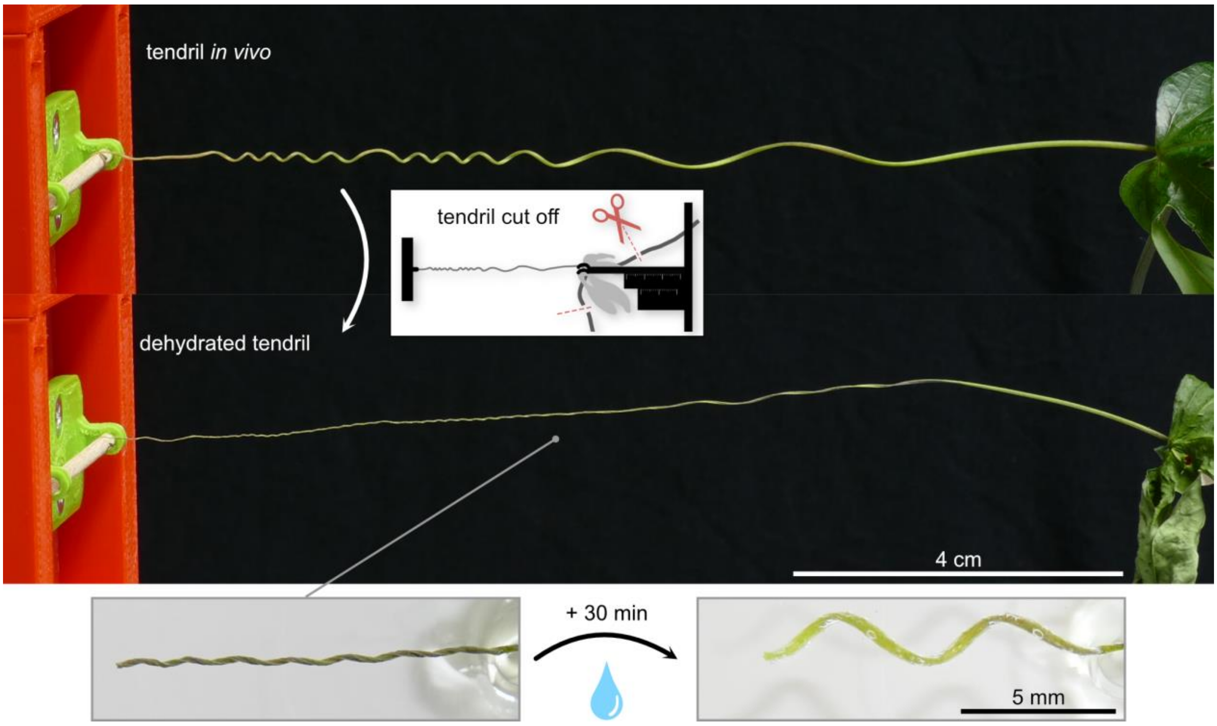


**Fig. S2: Rehydration of a dehydrated *P. caerulea* tendril segment.** After the *in vivo* coiling force generated by a *P. caerulea* tendril had been measured for 14.2 days, the plant stem was cut from the tendril and was left in the measurement setup to dry for 4.8 days (cf. fig. 3). When a segment from the dried tendril was submerged in tap water for 30 min, it gained a shape similar to that before the tendril was cut from the plant stem.

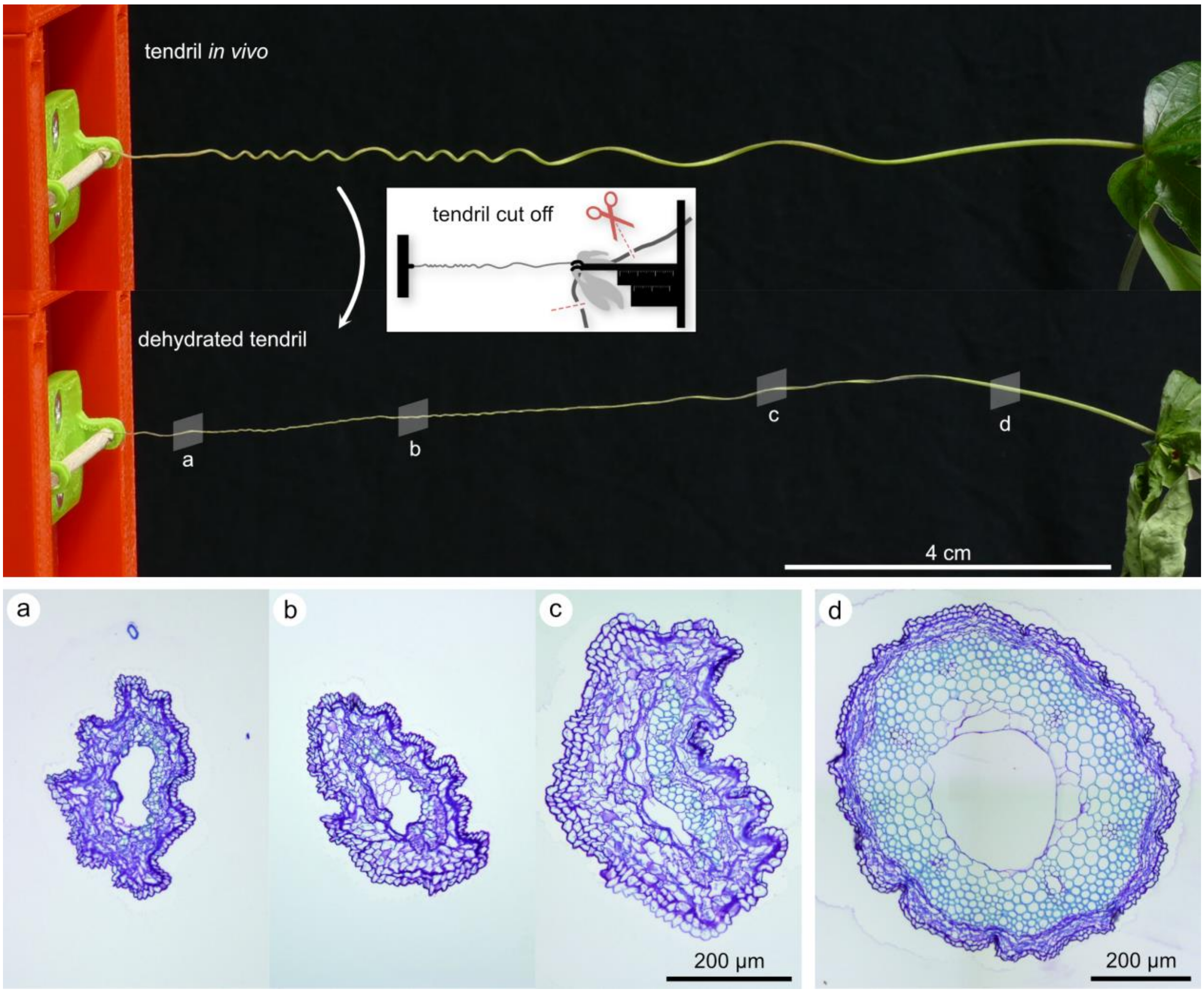


**Fig. S3: Tendril anatomy of a dehydrated *P. caerulea* tendril.** After measurement of the *in vivo* coiling force generated by a *P. caerulea* tendril for 14.2 days, the plant stem was cut from the tendril and was left in the measurement setup to dry for 4.8 days (cf. fig. 3). Cross-sections taken within the dried coiled part of the tendril **(a-c)** show that a large region of the tendril tissues collapsed, whereas in the straight basal part **(d)**, only epidermis, cortex, and phloem collapsed. Toluidine blue stained thin-sections.
